# When do measured representational distances reflect the neural representational geometry?

**DOI:** 10.1101/2024.12.30.630743

**Authors:** Veronica Bossio Botero, Nikolaus Kriegeskorte

## Abstract

The representational geometry of a brain region can be characterized by the distances among neural activity patterns for a set of experimental conditions. Researchers routinely estimate representational distances from brain-activity measurements that either sparsely sample the underlying neural population (e.g. neural recordings) or pool across the activity of many neurons (e.g. fMRI voxels). Here we use theory and simulations to clarify under what circumstances representational distances estimated from brain-activity measurements reflect the representational geometry of the underlying neural population, and what distortions must be expected under other circumstances. We demonstrate that the estimated representational distances are undistorted if single neurons are sampled at random. For voxels that take non-negatively weighted linear combinations, the resulting geometry is linearly distorted, correctly reflecting the population-mean dimension, while downscaling all orthogonal dimensions, for which the averaging cancels a large portion of the signal. Surprisingly, removing the mean from voxel patterns recovers the underlying representational geometry exactly in expectation under idealized conditions. This explains why the correlation distance, the most popular measure of representational dissimilarity in neuroimaging studies, “works” so well, yielding geometries that can appear similar between fMRI and neural recordings. The Euclidean (or Mahalanobis) distance computed after removing the mean of each pattern (without normalizing its variance) is an attractive alternative to the correlation distance in that it corrects for the inflated relative contribution of the population-mean dimension, while avoiding the drawback of the correlation distance: it can be large for confusable low-norm patterns, failing to reflect decodability. Our results demonstrate that measured representational distances reflect the neural representational geometry when (1) single neurons are sampled at random or (2) the weights with which the measured responses sample the neurons are drawn i.i.d. and (2a) the weights are drawn from a zero-mean distribution or (2b) the population mean is the same for all conditions or (2c) the mean is removed from each estimated pattern. We discuss practical implications for analyses of neural representational geometries.

## 1 Introduction

The representational geometry of a neural population code [29, 6, 32, 4, 22, 38, 27, 44] is characterized by the distances among the patterns of neural responses to a set of experimental conditions. Studying representational geometries enables researchers to abstract from the tuning of individual neurons and focus on which representational distinctions are emphasized and which are deemphasized, and to what extent, by the neural population as a whole. The representational distance matrix as a summary statistic enables direct comparisons between representations in brains and models, without requiring a correspondency mapping between the two representations [[30, 37, 43]; for related approaches, see also [24, 12]].

In practice, researchers estimate representational geometries based on experimental measurements of brain activity. However, the activity measurements are often severely limited samples of the underlying neural population activity in the brain region of interest. For example, single-cell recordings typically sample a small fraction of the neurons in the population. A study might sample 100 out of many millions of neurons in the cortical area studied. Functional magnetic resonance imaging (fMRI) achieves continuous coverage of the brain, but fMRI voxels (ranging from 4 mm in width to below 1 mm) reflect the average activity of tens or hundreds of thousands of neurons over several seconds[34].

Is it even possible to estimate the neural representational geometry from neural population activity measurements that subsample a tiny fraction of the dimensions? And if so, under what conditions will the representational distances we estimate from our data be good estimates of the true representational distances we would have estimated if we had recordings for the entire neural population?

### Surprising sensitivity

Representational geometries estimated from single-cell recordings and fMRI can appear strikingly similar. For example, Kriegeskorte et al. [31] compared inferior temporal representations of object images between macaques measured with sequential electrode recordings [674 cells, [23]] and humans measured with blood-oxygen-level-dependent (BOLD) fMRI using 2-mm isotropic voxels. The representational dissimilarity matrices for 92 object images were computed using the correlation distance (1 *— r*, where *r* is the Pearson correlation coefficient) and showed the same prominent categorical clusters and also significant within-category correlations of the representational dissimilarities between the macaque recordings and the human fMRI data. Other studies have found surprising sensitivity of BOLD fMRI to information thought to be inaccessible with voxels averaging across tens or hundreds of thousands of neurons. For example, the orientation of a visual grating can be decoded from the fMRI pattern it elicits in human primary visual cortex using 3-mm isotropic voxels [21, 17]. There has been some controversy as to whether (a) the 3-mm isotropic fMRI voxel patterns reflect subvoxel-scale neural pattern information through biased sampling or aliasing [21, 25, 1, 41, 2] or (b) the decodable information actually resides lower spatial-frequency bands of the neural activity patterns [39, 40, 14, 15]. This controversy about the sensititivity of BOLD fMRI with 3-mm voxels notwithstanding, there is evidence that fMRI using smaller voxels down to the submillimeter scale and ultra-high fields [8] can reflect columnar [47, 48] and even laminar [19] patterns of cortical activity.

### Voxels average locally, canceling contrast signals

Even if fMRI is surprisingly sensitive to fine-grained neural activity patterns, its sensitivity certainly falls off at the submillimeter scale. An undistorted reflection of the neural representational geometry would require that it is *equally* sensitive to all dimensions of the multivariate neural response space. Simulations show that the local averaging of activity in fMRI voxels can strongly affect the apparent representational geometry when the code is spatially structured at multiple spatial scales [26]. For a V1-like representation, for example, with local orientation columns and global retinotopy, orientation signals (which are carried by fine-grained patterns) will cancel more than location signals (which are carried by coarse-scale patterns). The more severe canceling of orientation than of location contrast signals distorts the apparent representational geometry. Kriegeskorte and Diedrichsen [26] concluded that accounting for local averaging in voxels is important to ensure correct inference of the data-generating brain-computational model. Empirically, it has been shown that face-identity information in the macaque face patch AL is prominently reflected in neural recordings but not fMRI data [13]. Some empirical fMRI studies have explicitly considered and modeled the effects of local averaging in voxels [42, 26, 5, 43].

The aim of this paper is to clarify under what conditions, if any, the neural representational geometry is correctly reflected in representational distance estimates that are based on measurements, as provided by cell recordings and fMRI, of only a small sample of the dimensions of neural population activity. By “correctly reflected”, we mean that our distance estimates reflect all aspects of neuronal tuning (e.g. orientation and position information for V1) equally, such that, if we had a sufficiently large number of measurement channels (e.g. recorded neurons or fMRI voxels), our distance estimates would be proportional (except for the effects of measurement noise) to the neural representational distances we would have obtained from recordings of the entire neural population.

We will describe the ideal conditions under which the estimated distances are undistorted in expectation. Note that under such ideal conditions, the following two challenges for analysis remain: (1) There would still be random variation of the estimates arising from noise in the measurements and from the random sampling of a limited number of measurement channels. (2) Naive estimation of the representational distances with non-negative estimators would still suffer from positive estimation bias. These two issues have been addressed in previous work on unbiased distance estimation [28, 37, 45, 11, 27] and statistical inference on representational geometries [37, 43] and are orthogonal to the question we are concerned with here.

### Cell recordings and fMRI sample projections of the geometry

We can think of both electrodes and voxels as sampling *projections* of the neural representational geometry. Imagine measuring the activity of a population of *N* neurons as they respond to a set of stimuli. Each response pattern of the population can be viewed as a point in and *N* -dimensional space. We can think of each neuron’s activity as encoding the coordinate along a particular axis of this space (where the axes are by definition orthogonal). The set of unique response patterns to *P* stimuli constitutes a point cloud in this *N* -dimensional space, whose pairwise distances define the neural representational geometry (Fig. 1a). Taking measurements of these neurons can be viewed as projecting the point cloud onto a new set of axes determined by the measurement channels (Fig. 1b). For example, an electrophysiological recording of individual neurons with an electrode array can be viewed as a projection of the point cloud onto the subspace spanned by the axes corresponding to the neurons captured by the array. In the example shown in Fig. 1a, the entire population consists of three neurons and the measurement of the first two neurons would result in an orthogonal projection of the activity patterns onto two axes defined by the measurement channels (Fig. 1b).

**Figure 1:**
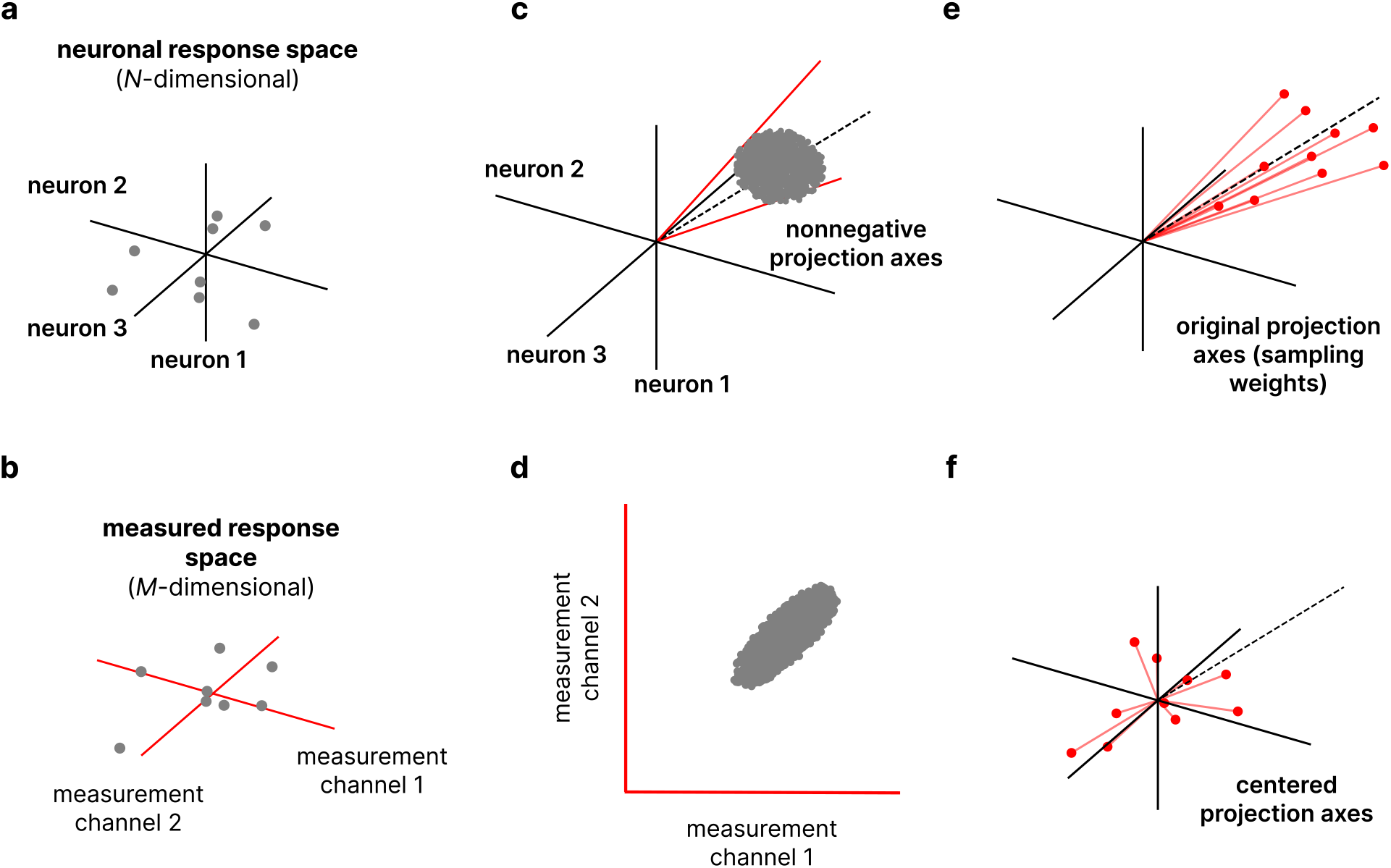
Brain activity measurements as sampling projections of the neural response patterns. a. Neural activity response patterns can be viewed as points in a multidimensional metric space in which the dimensions correspond to the activity of single neurons. **a.** Example of a population of three neurons, each gray point represents the population response to one experimental condition. **b.** Projection of the neural response patterns onto the measured response space. Each point is recoded along a new set of axes given by the measurement channels. **c.** Measurement channels that reflect the activity of the neural population as weighted averages with only non-negative weights can be viewed as sampling axes that lie on the all-positive orthant. Panel **c** shows a set of response patterns with a spherical geometry (gray) and two example all-positive sampling axes (red). **d.** Recoding the points on the sphere along the sampling axes results in a geometry that is linearly stretched along the dimension of the neural population average. **e.** A set of non-negative sampling axes. Each red vector represents a measurement channel whose entries correspond to the neural sampling weights. These vectors can also be seen as points in the neural response space, represented by the red tips. The fact that all the weights are non-negative makes the sampling axes closely aligned with the all-1 vector.. **f.** Removing the mean activation from each measured pattern is equivalent to rigidly centering the tips of the original sampling weight vectors on the origin of the neural response space. This corrects the oversampling of the population-mean (all-1) axis, dispersing the effective axes along which the neural patterns are sampled, such that all directions are equally represented in expectation and the neural representational geometry is thus undistorted.

A representational distance is a property of the neural population. Estimating it from a sample of neurons is analogous to estimating other population parameters from samples. Consider estimating the height of the Chilean population by measuring the height of a random sample of 500 Chilean individuals. We would expect the average height of the random sample to be a good estimate of the average height of the Chilean population. In estimating the representational distance, we would expect the average squared difference across a sufficiently large random sample of particular neurons to be a good estimate of the average squared difference across the entire neural population. The key thing to note in this analogy is that representational distances can be estimated as averages of the squared differences between neural activities. The sample average of the squared differences is a good estimate of the population average of the squared differences, and the latter is proportional to the population squared Euclidean distance. The proportionality constant depends on the number of neurons. However, researchers using representational similarity analysis [30, 37, 9, 43] usually disregard the scaling and rely only on the distance ratios to make inferences.

If, instead of sampling a subset of neurons, each channel measured a linear combination of the activity of the neurons in the population with random weights, the measurement process would amount to a projection of the point cloud onto a set of randomly oriented vectors [16]. Random projections can yield undistorted distance estimates [20]. In particular, if the projection weights are drawn i.i.d. from a zero-mean distribution, then the distances will be undistorted in expectation. Moreover, the Johnson-Lindenstrauss Lemma [20] states that the number of measurement channels required for a good approximation does not depend on the number of neurons and grows only logarithmically with the number of stimuli. The Johnson-Lindenstrauss Lemma thus suggests (1) that even measurements that combine many neurons might yield undistorted estimates of the geometry, and (2) that a realistically modest number of measurement channels might suffice.

Unfortunately, the Johnson-Lindenstrauss Lemma doesn’t directly apply to either fMRI voxels or cell recordings, where the sampling weights will not have expected values of zero. We can think of fMRI voxels as taking locally extended and overlapping non-negatively weighted averages [25] and of array recordings as taking sparse punctate samples. In either case, the linear combinations will not use negative weights, and thus the mean weight will not be zero.

### Contribution of this paper

Using theoretical analysis and simulations we demonstrate several facts. If each sample consists of a single neuron and neurons are sampled at random, the estimated representational distances are undistorted and even just 50 neurons can yield a good correlation with the ground-truth distances. Sampling neurons with (hypothetical or simulated) voxels that use non-negatively weighted linear combinations, where the weight of each neuron is drawn i.i.d. from a non-negative weight distribution, results in a geometry that correctly reflects the population-mean dimension, while deemphasizing dimensions orthogonal to the population-mean dimension. The resulting apparent geometry is scaled down uniformly for all dimensions orthogonal to the neural-population mean dimension, while remaining unscaled along the neural-population mean dimension. Because we always disregard the overall scaling when comparing representational distance matrices, we can equivalently (and perhaps more intuitively) imagine the apparent geometry as linearly stretched along the neural-population-mean axis.

Pattern information analyses and representational similarity analyses often remove the regional-mean activity from each pattern before assessing the distance or decoding accuracy [28, 35]. This normalization step can be motivated as making pattern analyses more complementary to regional-mean activation analyses. An alternative motivation is that removing the mean from each pattern deemphasizes the neural-population mean axis, which is overemphasized in fMRI relatively to all other dimensions of the neural population response. Removal of each pattern’s mean is also implicit to the correlation distance, which is the most widely used representational dissimilarity measure.

Removing the mean from each fMRI pattern estimate may be expected to reduce the information content. However, the mathematical and simulation results of this study show that it can help reveal the underlying neural representational geometry, correcting the distortion caused by local averaging if voxels sample random sets of neurons. Our results explain why the correlation distance “works” so well as a measure of representational dissimilarity in the context of fMRI data. [31] used the correlation distance to measure representational dissimilarities and revealed a close match between the representational geometries estimated from human fMRI and monkey cell recordings. We show below how the mean removal implicit to the correlation distance revealed the matching representational geometries.

The correlation distance has been by far the most popular representational dissimilarity estimator in fMRI [33, 18, 36, 29, 46, 7]. The correlation distance is proportional to the squared Euclidean distance measured after removing the mean and normalizing the variance of each pattern (see Supplemental material 7.4). The variance normalization causes a serious drawback: The correlation distance cannot be interpreted in terms of stimulus decodability because identical patterns of low amplitude will have low correlations (because noise dominates) and therefore large correlation distances; [45]. Our results here suggest that the benefits of the correlation distance result from mean removal. Removing the mean from each pattern is well motivated by the goal to investigate the underlying neural representational geometry – within the limits imposed by fMRI. We can get the benefit of mean removal without the drawback of variance normalization, by replacing the correlation distance by the squared Euclidean or Mahalanobis distance (or an unbiased estimator of these; [9]), computed after mean removal.

The structure of the paper is as follows. We begin by formulating the problem of estimating representational geometries from measurements of brain activity in mathematical terms. Our theoretical analysis uses models of neural activity sampling to reveal the distortion of the neural representational geometry that is expected. These analyses reveal, in particular, under what circumstances distance estimates are guaranteed to be undistorted in expectation despite limited measurements. We then describe simulations based on ground-truth geometries to confirm the theoretical results and illustrate the effect that removing the mean from the measurement patterns has in correcting the distortion. Finally, we discuss the implications of these results for interpreting past empirical findings and for designing future analyses of representational geometries.

## 2 Results

### 2.1 Theory

We consider measuring the responses of *N* neurons with *M* measurement channels. Each measurement channel *π_i_* (*i* = 1*..M*) reflects the neural activity pattern **x** (*N ⇥* 1 column vector) as a linear combination with random weights *w_ij_* ∈ ℝ drawn i.i.d. from some probability distribution over the real numbers (*j* = 1*..N*). The univariate distribution from which each *w_ij_* is independently drawn could be of any shape. In particular, it could be a zero-mean Gaussian, a Gaussian with a positive mean, a non-negative distribution, or a binary distribution with a spike at zero and another at one. We can think of fMRI voxels as sampling with weights drawn from a non-negative distribution: Each voxel takes a local non-negatively weighted linear combination of the neural activity. We can think of invasive neurophysiological recordings as sampling with weights from a binary distribution with a spike at zero and another spike at one: Each electrode tip takes a punctate sample of the neural activity, perhaps reflecting spiking activity of multiple local neurons.

We assume here that the measurements are not affected by noise. Noise creates a positive bias for distance estimates, which can be removed through cross-validation [45]. However, this is an orthogonal issue to the one we consider here. The bias we are concerned with is not caused by noise, but by the greater sensitivity to the neural population-mean dimension of the neural response space that arises when measurement channels average with non-negative weights across the neurons.

#### 2.1.1 Measured dissimilarities are unbiased for zero-mean neural sampling weights or equal-mean neural patterns

For this independent sampling model, the expectation of the squared Euclidean distance between the measurements *π*(**x**) and *π*(**y**) (*M* × 1 column vectors) of two neural activity patterns **x** and **y** decomposes into two additive terms:

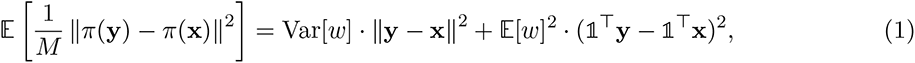

where

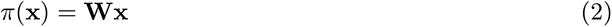

**W** is an *M* × *N* matrix whose entries are *w_ij_*, drawn i.i.d. from a univariate weight density. (Proof in Supplemental material 7.1. See also, [43])

Equation 1 shows that the neural population mean dimension of the neural response space is overrepresented relative to all other dimensions when the representational dissimilarities are naively computed from the measurement channels, unless the univariate weight distribution has mean zero 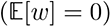. All other dimensions are represented without bias as the measurements take random linear combinations of neural activity. If either the mean of the weight distribution is zero or the mean neural activity is the same for each activity pattern 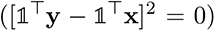 then the second term in Eq. 1 is zero, and all representational dissimilarities are correctly reflected in the measured representational dissimilarities in expectation.

#### 2.1.2 The apparent representational geometry is linearly stretched along the all-1 direction of the neural response space

For the linear random sampling model we consider here, the apparent representational geometry (i.e. the geometry implied by the Euclidean distances among the measured patterns) is distorted in a particularly simple way: It is stretched along the all-1 direction of the neuronal response space (Fig. 1 c, d). The assumption that each measurement channel randomly samples the neurons entails that all directions orthogonal to the all-1 direction are equally represented. However, unless the weight distribution has a mean of 0, the projection axes corresponding to the measurement channels (Fig. 1 c, e) are somewhat aligned with the all-1 vector, oversampling the component along the all-1 direction, which causes the stretching of the apparent representational geometry along the all-1 direction.

Note that we care only about the relative representation of components along the all-1 direction and orthogonal to this direction, not about the overall scale factor. This justifies thinking of the geometry as *stretched* along the all-1 direction. If we cared about the absolute distances, it would be more appropriate to think of the geometry as linearly shrunk along all directions orthogonal to the all-1 vector because it is signal cancellation caused by averaging that reduces our sensitivity to pattern differences orthogonal to the all-1 vector, whereas averaging neural responses does not reduce our sensitivity to changes of the mean of the neural responses.

The factor *f_shrink_* by which orthogonal directions are shrunk relative to the all-1 direction follows directly from Eq. 1 (see Supplemental material 7.2 for derivation):

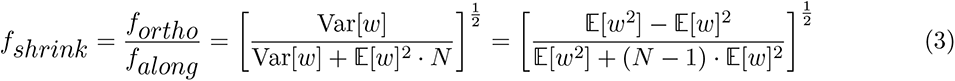

To see how this factor follows from Eq. 1, consider that the second term of Eq. 1 is zero for two neural response patterns differing only orthogonal to the all-1 vector of the neural response space, so the measured squared Euclidean distance is the neural squared Euclidean distance times Var[*w*]. For two neural response patterns differing only along the all-1 vector in the neural response space, by contrast, both terms from Eq. 1 matter, and the measured squared Euclidean distance is the neural squared Euclidean distance times Var[*w*]. Note that, consistent with the discussion above, the geometry is undistorted when the weights distribution has expected value 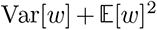, which renders the ratio 1.

#### 2.1.3 Dissimilarities among the mean-removed measured patterns provide unbiased estimates of neural dissimilarities

We now show a perhaps counterintuitive fact: When we remove the mean activation across channels from each measured activity pattern, then the squared Euclidean distances among the (mean-removed) measured patterns correctly reflect the neural dissimilarities without any bias in the representation of the neural population mean dimension.

It is intuitive that removing the mean of each measured pattern would reduce the overrepresentation of the neural population mean dimension that results when the weights have non-zero mean. However, it is not obvious why removing the mean of each measured pattern *exactly* removes the bias and results in dissimilarity estimates that, in expectation, correctly reflect both the neural population mean dimension and all other dimensions, without either over- or underrepresenting the neural population mean dimension.

If the mean of the measurements perfectly reflected the neural population mean, then removing the mean of each measured pattern would collapse all variability along the neural population mean and overcorrect: Any differences along the neural population mean dimension would not be reflected in the measured representational dissimilarities at all anymore. However, the mean of the measurements does not perfectly represent the neural population mean dimension. The measurement mean deviates in such a way that removing it entails appropriate representation of the neural population mean dimension relative to all other dimensions.

Removing the mean from the measured voxel pattern **v** = **Wx** to obtain the centered voxel pattern v̅ can be achieved by pre-multiplication with a centering matrix 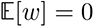, which centers each incoming column:

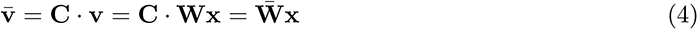

Centering the voxel pattern **v** is equivalent to centering each column of the neural sampling weights matrix **W**. The centered voxel pattern v̅, thus, behaves as if it were an activity pattern measured with zero-mean neural sampling weights 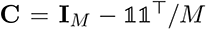, where the bias (second term of right hand side in Eq. 1) vanishes (see Supplemental material7.3.1 for more details).

Note that centering the measured patterns (or equivalently each column of **W**) is not exactly equivalent to drawing weights i.i.d. from a centered version of the univariate distribution that the weights are drawn from. If the univariate weight distribution were centered, then each column of **W** would not exactly sum to 0. The expected value of the sum of weights would be 0, but there would be a small deviation reflecting the randomness of the sample. However, in expectation, centering the measured patterns yields the same distance estimate as measuring with a technique where each “voxel” samples with 0-mean weights.

To understand the effects of removing the mean from each pattern intuitively, we need to consider the *rows* of **W**, the neural sampling weight vectors, whose tips we can think of as an ensemble of points in the (high-dimensional) neural response space (Fig. 1e). Centering each column of **W** amounts to centering of the ensemble of tips of the neural sampling weight vectors on the origin of the neural response space (Fig. 1f). As the tips are moved together (each along the same vector, thus rigidly maintaining their geometric relationships), each vector continues to emanate from the origin, so the direction it points in changes. The changes of direction entailed by centering the ensemble of tips changes the dimension of neural response space that each vector samples. The projection axes are then no longer aligned around the all-1 vector.

Note that centering the ensemble of tips does not change the geometry of the ensemble (the shape of the distribution of tips) over the neural response space. It only changes the means (making the mean weight for each neuron zero). As a consequence, projecting the weight vectors onto any given axis of neuronal response space yields the same (scalar) variance when done before or after centering. This is why the centering yields an equal distance estimate for the measurement of unit-norm pattern difference vectors of any orientation in expectation. After centering, the projection axes equally sample all directions.

The formal result that the squared Euclidean distance between the mean-removed measurement patterns 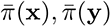 provides an unbiased estimate of the distance between the original neural patterns can be stated as follows:

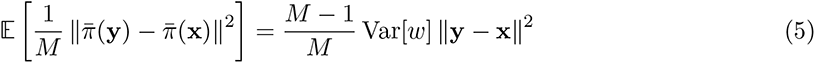

(See Supplemental material 7.3.2 for proof.)

### 2.2 Simulations

#### 2.2.1 Simulations based on random ground-truth geometries

We simulated neural activity patterns for a set of experimental conditions by drawing points randomly from a multivariate probability distribution. This provided a ground-truth representational geometry that we compared to the geometries obtained by measuring the activity patterns with different sampling models. We used sampling models that mimic the effect of measuring a population of neurons using single-cell recordings and fMRI.

##### Simulations confirm theoretical results

Fig. 2 shows representational dissimilarity matrices (RDM) for an example ground-truth simulated geometry and the geometries obtained by sampling the simulated neural patterns with linear combinations of the neurons. We consider two variants of an independent-weight-linear-combination-sampling (IWLCS) model. In the random-projection sampling model, weights are drawn i.i.d. from a zero-mean Gaussian. In the voxel sampling model, weights are drawn i.i.d. from a non-negative distribution. In the voxel model, thus, each channel reflects a non-negatively weighted linear combination of the neuronal activity (see Methods, 4.1.2). The dissimilarity matrices illustrate that the distances estimated using the raw measurements from the voxel sampling model (Fig. 2, bottom middle) are poorly correlated with the true neural distances (Fig. 2, top left). The distorted distance estimates are highly correlated with the absolute differences of the population-mean response (Fig. 2, bottom left), illustrating that the population-mean dimension is overrepresented in the measurements. As predicted by the theoretical results, removing the mean from the measured patterns before estimating the dissimilarities visibly corrects for the distortion, yielding an RDM that captures the ground-truth distances (Fig. 2, bottom right). For the random projection reference model, both the raw pattern distance estimates (top middle) and the mean-removed pattern distance estimates (top right) reflect the ground-truth geometry well. The mean removal here is similar to removing a randomly chosen dimension and does not strongly alter the apparent geometry.

**Figure 2:**
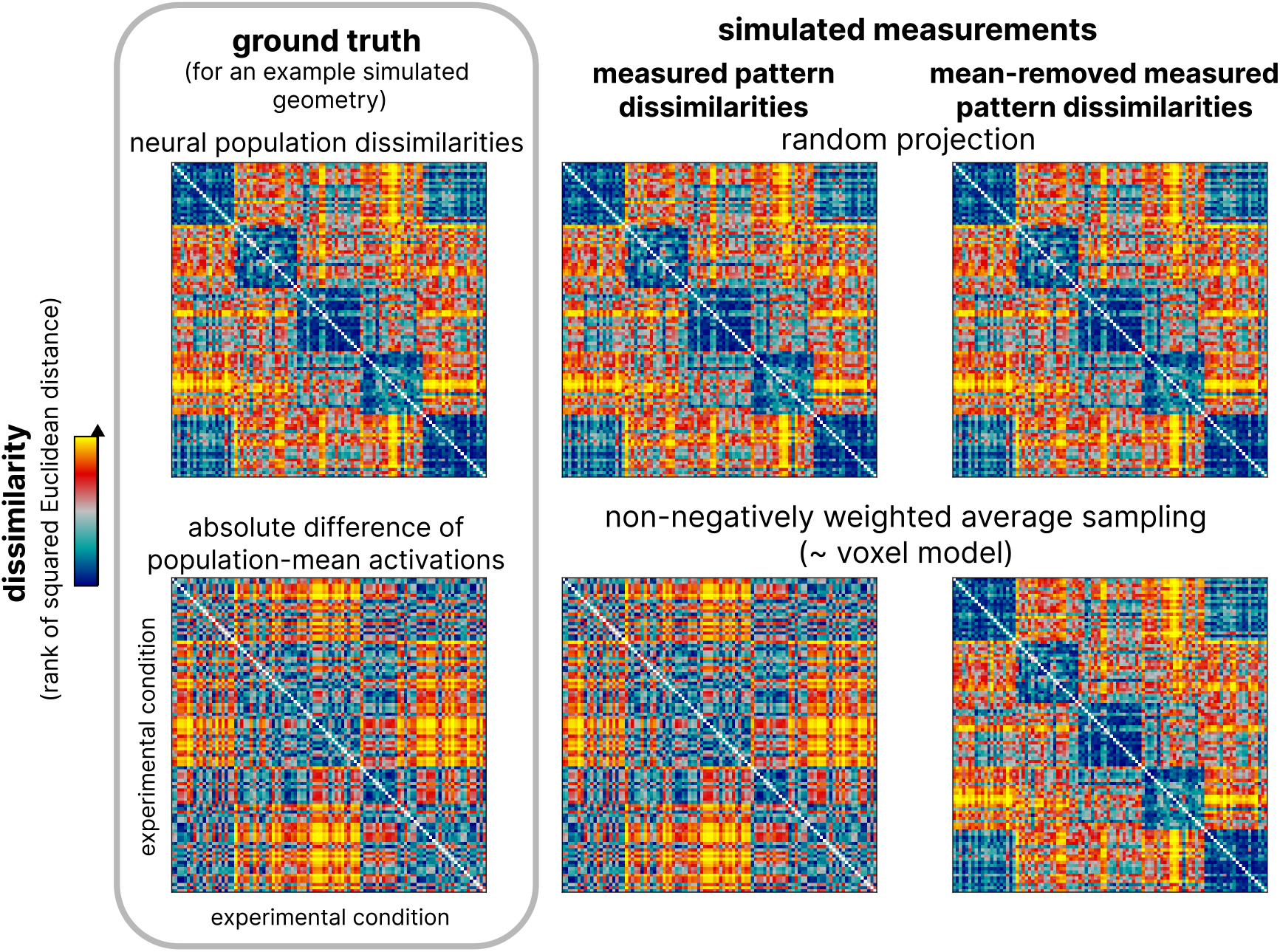
Representational dissimilarity matrices for an example randomly generated geometry and simulated measurements. The left column shows squared Euclidean distances between the pairs of simulated neuronal patterns (top) and the absolute difference after averaging over all the neurons for each activity pattern (bottom). The middle column shows the distances between the patterns obtained after a random projection (top) and a simulation of voxel sampling (bottom). The right column shows the squared Euclidean distances between the pairs of activity patterns after subtracting the mean over the channels for each individual simulated experimental condition for random projection sampling (top) and simulated voxel sampling (bottom). For random projections (top), note that mean removal amounts to the removal of one of many dimensions of the representational space and does not appreciably alter the apparent representational geometry. For the voxel sampling model (bottom), mean removal corrects the overemphasis on the population-mean dimension (which results from averaging within voxels), revealing the otherwise hidden neural representational geometry.

We quantified these results by calculating the Pearson correlation coefficient between the simulated ground-truth RDMs and the measured RDMs, varying the number of measurement channels. For each simulation, we randomly generated a new ground-truth geometry. The results across simulations are shown in Fig. 3. We simulated the measurement process using two models of fMRI voxel sampling (both non-negative IWLCS models, one of them sparse), one model of single-cell array recordings (using a subsample of the neurons), and random projections for reference (for details on the measurement models, see Methods, section 4.1.2).

**Figure 3:**
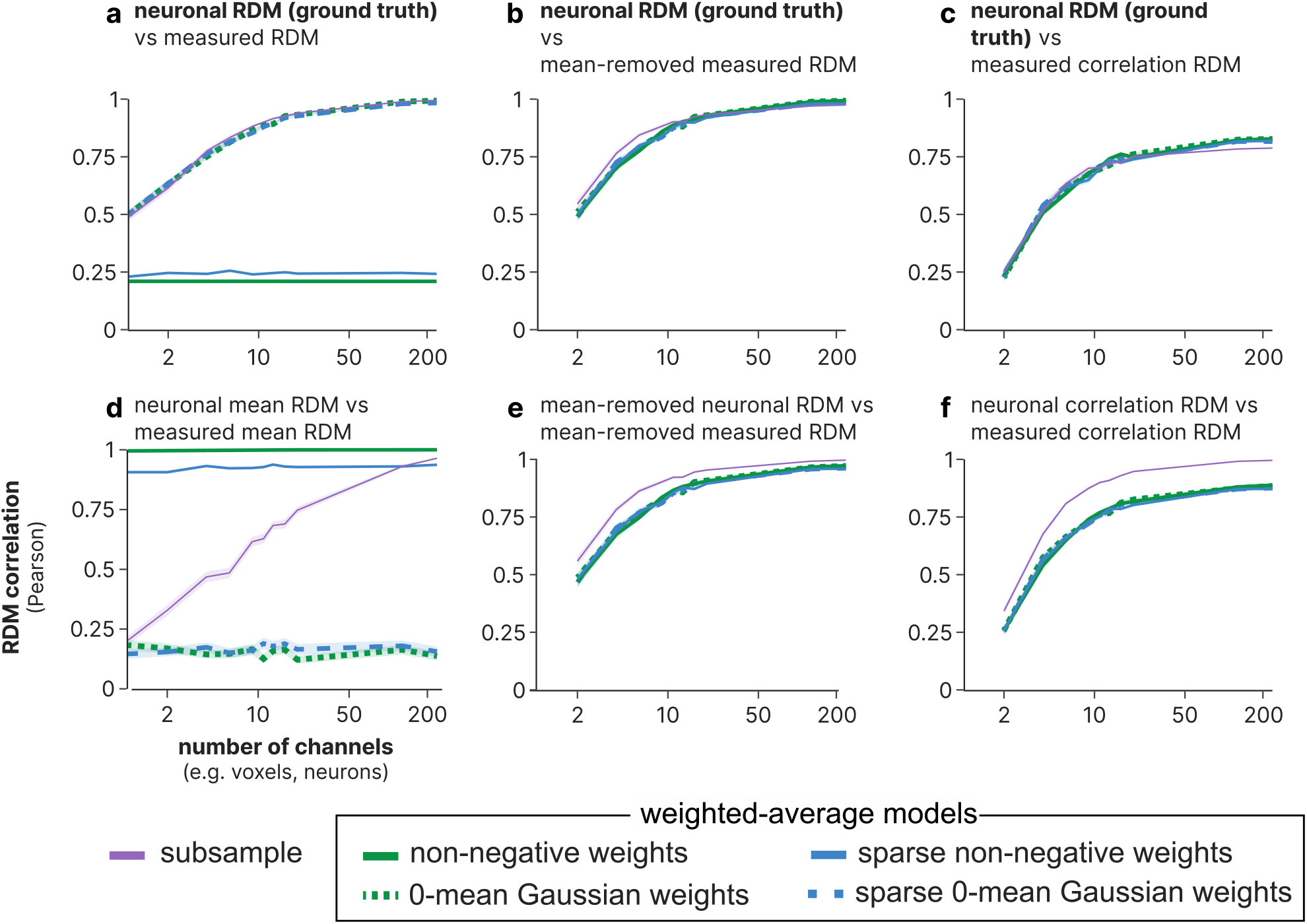
Correlation between representational distances of ground-truth simulated activity patterns and simulated measured activity patterns. Plots show the Pearson RDM correlation (vertical axes) between the simulated neural pattern RDM and the simulated measured pattern RDMs for five measurement models (lines) averaged across simulations. The shaded region around each individual line indicates the standard error of the mean across simulations. For each of a sample of randomly generated representational geometries, we simulate measuring the neuronal population varying the number of channels (horizontal axis). Panels show the RDM correlation between: **a.** the squared Euclidean distance RDM of the ground-truth simulated neuronal patterns and each of the squared Euclidean distance RDMs for simulated measured patterns. **b.** the squared Euclidean distance RDM of simulated neuronal patterns and the RDM of simulated measurement patterns after mean removal. **c.** the squared Euclidean distance RDM of the simulated neuronal patterns and the correlation distance RDM of the simulated measurement patterns. **d.** the RDMs obtained by taking the absolute differences of the average over all the simulated neurons/channels for each activity pattern. **e.** the RDM of simulated neuronal patterns after mean removal and the RDM of simulated measured patterns after mean removal. **f.** the correlation distance RDM of the simulated neuronal patterns and the correlation distance RDM of the simulated measurement patterns.

The neural distances are captured equally well by the random projection models and the subsample model (Fig. 3a). In contrast, non-negative IWLCS models result in distances that correlate poorly with the neural distances regardless of the number of measurement channels (Fig. 3a). Notably, the mean component of the ground-truth RDM is recovered accurately from the population-mean distances obtained with the non-negative weights models, which illustrates the overrepresentation of the population-mean dimension resulting from non-negative sampling (Fig. 3d). Consistent with the theoretical results, removing the mean from the measured patterns before estimating the dissimilarities recovers the underlying geometry in expectation. The mean-removed RDM is highly correlated with the ground-truth RDM for every model for as few as 50 measurement channels (Fig. 3b).

The theoretical results, perhaps counterintuitively, suggested that removing the mean from the measured patterns does not entail undersampling of the true neural population mean. To assess whether this prediction held in the simulations, we removed the ground-truth population mean from the ground-truth neural patterns and compared the resulting simulated neural pattern dissimilarities to the mean-removed measured pattern dissimilarities. For the non-negatively weighted average models, this comparison yields lower RDM correlations than for the original ground-truth neural patterns, consistent with the prediction from the theoretical results that the mean-removed measured patterns reflect all dimensions (including the true neural population-mean dimension) equally in expectation (Fig. 3e).

##### Removing the mean from measured activity patterns provides better estimates of the neural representational geometry than using the correlation distance

A common practice when analyzing measurements of brain activity using RSA is to use the correlation distance to characterize the geometry of the measured patterns. The correlation-distance RDM is proportional to the squared Euclidean distance RDM computed after mean removal and variance normalization of each pattern (details in Supplemental material 7.4). Given the implicit removal of the mean across channels from each pattern when using the correlation RDM, we asked how well the correlation-distance RDM captures the simulated neuronal RDM. We found that using the correlation-distance RDM yields similar RDM correlations as using the squared-Euclidean RDM after mean-removal (without normalizing the pattern variance). The correlation distances among the measured patterns are highly correlated with the squared Euclidean distances in the ground truth geometry (Fig. 3c). As before, when we removed the ground-truth neural population-mean from the groundtruth neural patterns, the RDM correlation with the mean-removed measured pattern RDM was reduced (Fig. 3e, f).

Our results show that pattern mean removal alone (without the pattern variance normalization implicit to the correlation distance) yields significantly higher RDM correlations with the ground-truth neural RDM for both voxel models tested (Fig. 4). The lower RDM correlation with the ground-truth RDM can be explained by the additional transformation introduced by the correlation distance: the divisive normalization by the variance for each pattern. Our simulations suggest that this extra step degrades the accuracy of distance estimates.

**Figure 4:**
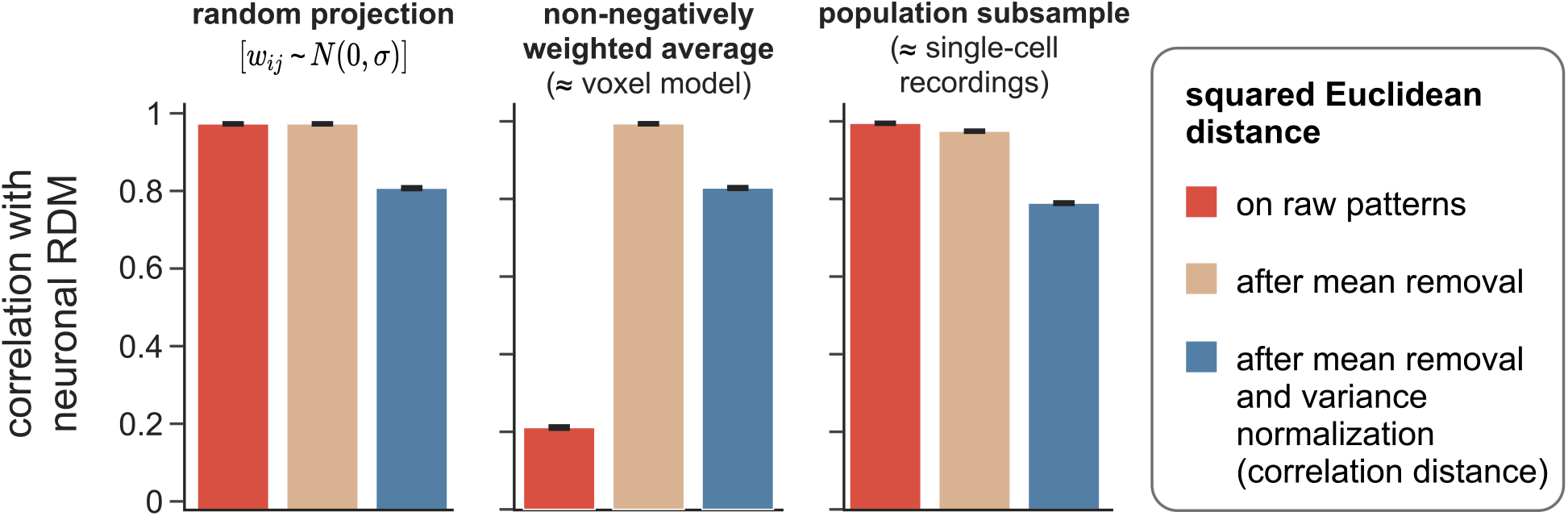
Pattern-mean removal alone is more effective than pattern-mean removal and pattern-variance normalization at correcting the population-mean overemphasis in the measured representational geometry. We can compute the squared Euclidean distance on raw (red) or normalized measured patterns. Normalization can consist in subtracting the mean across measurement channels (e.g. voxels) from each pattern (beige) or in subtracting the mean and applying divisive normalization to scale the pattern variance to 1 (blue). When the squared Euclidean distance is computed after mean and variance normalization, it equals twice the Pearson correlation distance (derivation in Supplement Section 7.4) and thus yields the same RDM correlations (blue). Bars indicate the RDM correlation between the ground-truth neural RDM (squared Euclidean distance) and the RDMs containing the squared Euclidean distances among the measured patterns. Each subplot corresponds to a a different measurement model. If the measurements sample random projections (weights drawn i.i.d. from a 0-mean Gaussian, an unrealistic scenario shown in the left panel), the squared Euclidean distances of the measurement channels reflect the ground-truth neuronal representational geometry well. Mean removal hardly aflects the RDM correlation, but mean removal and variance normalization (implicit in the correlation distance) performs markedly worse. If the measurements sample non-negatively weighted averages (a scenario comparable to fMRI voxels, center panel), mean removal greatly improves the RDM correlation with the ground-truth neuronal RDM in the estimated RDM. Additional normalization of the pattern variance (or, equivalently, the use of correlation distance) markedly degrades the estimate of the neural representational geometry. If the measurements consist in a random subsample of the neurons (an idealization of the typically somewhat biased samples in neural recording experiments, right panel), the estimate of the representational geometry is highly accurate. Mean removal then slightly degrades the RDM correlation with the ground truth. Mean removal and variance normalization markedly degrade the RDM correlation with the ground truth. The error bars represent the standard error of the mean across simulations with different ground-truth representational geometries

#### 2.2.2 Simulations based on neural data

[31] analyzed inferior temporal (IT) activity patterns elicited by images of objects belonging to different categories in humans and macaques. The activity patterns were measured using fMRI in humans (blood-oxygen-level-dependent fMRI at 3 Tesla using 2-mm-wide isotropic voxels) and cell recordings in macaques (data set from [23]). The representational dissimilarity matrices obtained in this study were highly correlated between the two species. We now use simulations based on these empirical data to provide a plausible explanation as to why the representational geometry appears to be conserved despite the radical difference in the two measurement techniques.

Using the empirically observed dissimilarities among stimuli in the monkey IT, we created embeddings of high-dimensional patterns corresponding to neural activations in response to the visual stimuli (see Methods Section 4.2). We used these patterns as the ground-truth reference geometry and, as before, sampled this simulated neural population using various measurement models.

Fig. 5 shows RDMs from one example measurement simulation with 50 channels. The squared Euclidean distances between patterns estimated from the voxel sampling model appear distorted relative to the ground-truth pattern distances. The RDM obtained after mean-removal, in contrast, visually closely resembles the neural RDM. For the subpopulation sampling model both RDMs apparently reflect the true geometry well.

**Figure 5:**
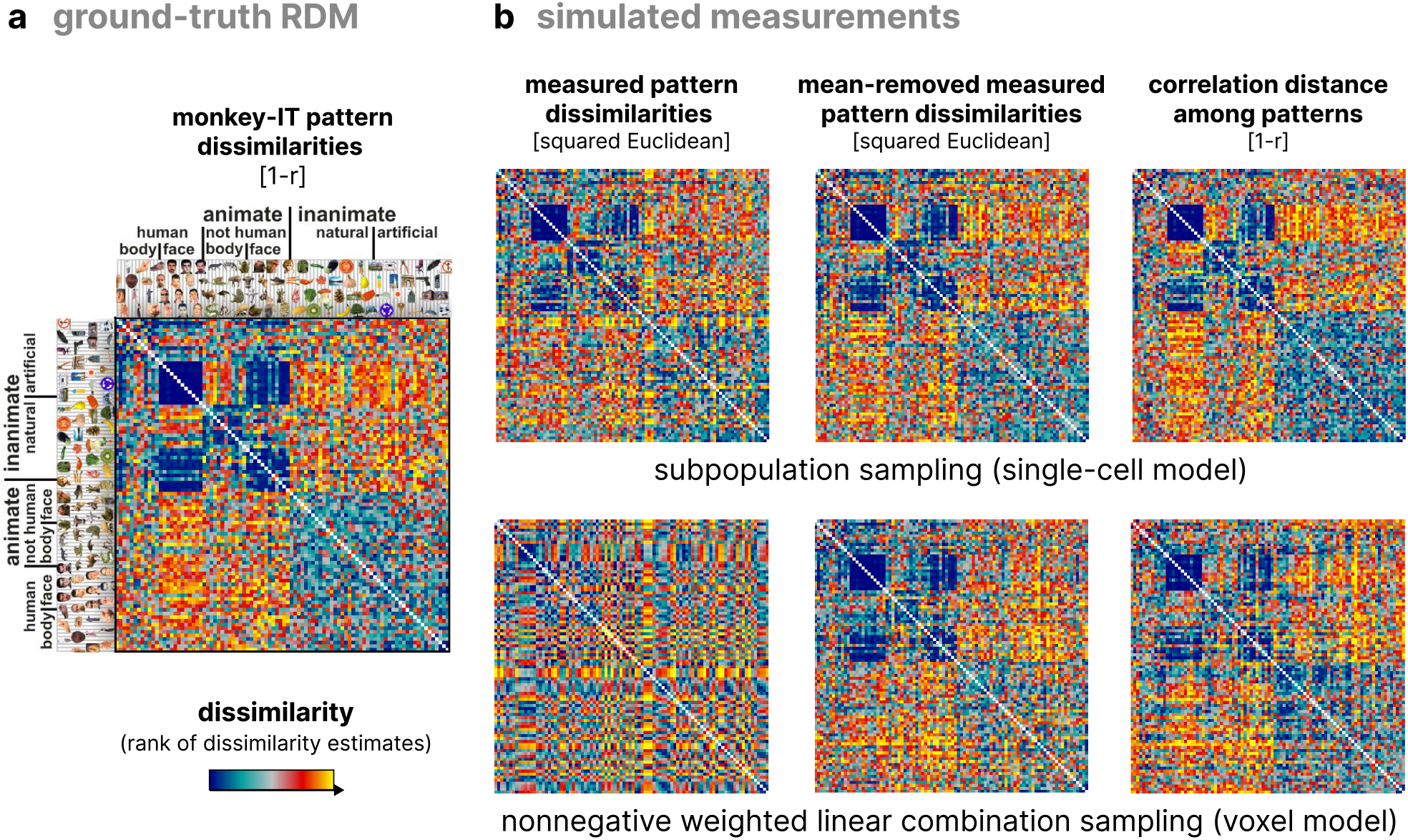
Representational dissimilarity matrices for empirical data and simulated measurement patterns. Panel **a** shows the RDM of the distances between the activity patterns generated from cell recordings of 674 neurons in monkey IT [23, 31], which we use as the ground-truth geometry for the simulated measurements. Panel **b**: Left column shows the RDM of squared Euclidean distances between simulated measurements taken with the subpopulation sampling model (top) and non-negative IWLCS model (bottom). Middle column shows the squared Euclidean distance RDMs taken from the simulated measurements after removing the mean from each measured pattern. The right column shows the RDMs obtained using the correlation distance on the simulated measurements, which implicitly removes the mean, and matches the analyses used in the empirical study.

We replicated the analyses from [31] on our simulated measurements. To match the study’s methods we used the Pearson correlation distance (1-r) as the dissimilarity estimator (Fig. 5, right). The correlation distance implicitly removes the mean from each measured pattern. The distribution of representational dissimilarities between ground-truth response patterns and experimental and simulated measurement patterns are shown in Fig. 6. The correlations obtained with the voxel sampling model (Fig. 6b) closely resemble those observed with the fMRI measurements from human IT in the original study (Fig. 6a). The subpopulation model yielded correlations that closely matched the monkey IT correlations (Fig. 6c).

**Figure 6:**
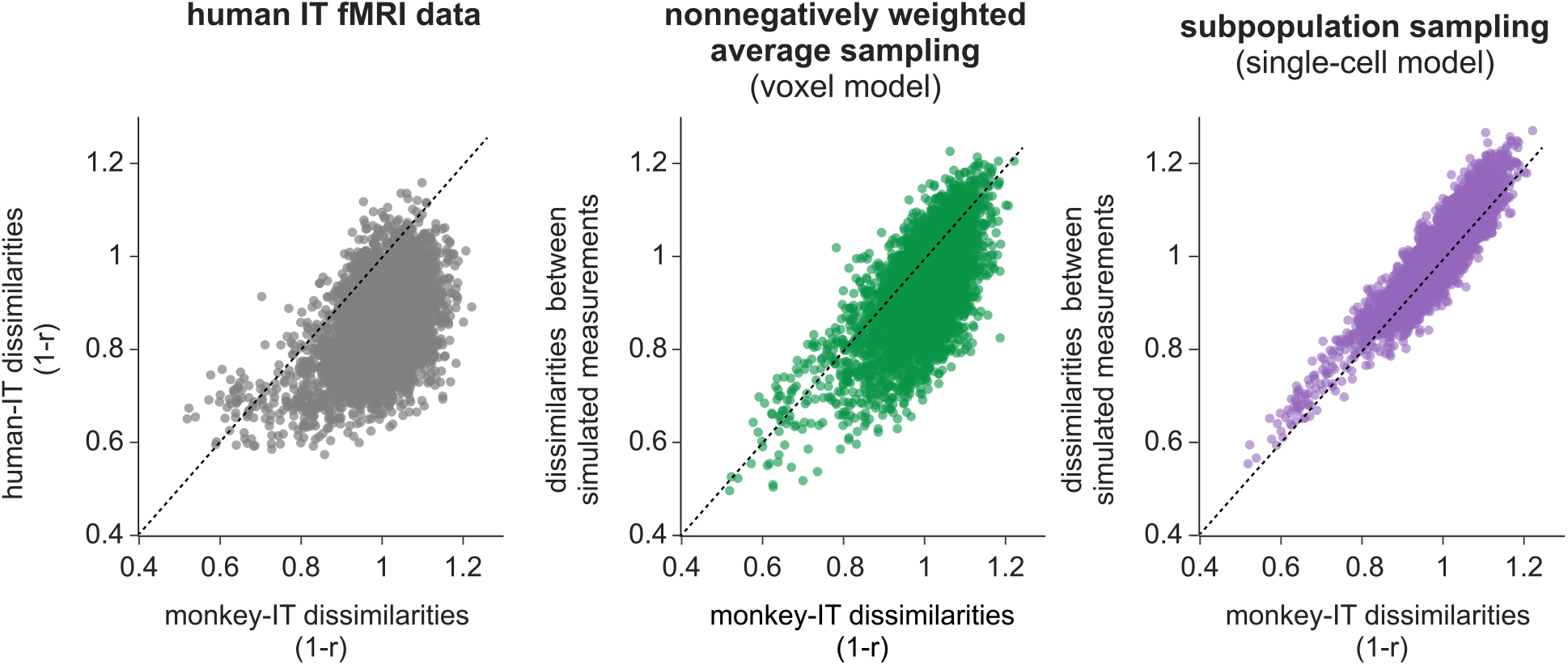
Correlation of representational dissimilarities between monkey and human IT for empirical data and simulated measurements. Panel **a** shows the results from the empirical data in the original study. Each dot marks the correlation between activity patterns for a pair of stimuli as measured in monkey IT (horizontal axis) and human IT (vertical axis). Simulation results are shown in panel **b**, where voxels were simulated as sampling a neural population with non-negatively weighted means. The simulated neural population encoded a representational geometry as observed in the monkey data, so the neural pattern dissimilarities (correlation distance: 1 *— r*, horizontal axes) match those of the actual neural recording data (left panel). These ground-truth dissimilarities are plotted against the dissimilarities among thesimulated voxel response patterns (correlation distance: 1 *— r*, vertical axis). Panel **c** shows results of a similar simulation, where the measurement model selects a random subset of neurons (subpopulation sampling). The concentration around the diagonal shows that the representational dissimilarities (1*—r* as before for both sampling) can be quite accurately estimated from a subpopulation of 200 neurons, if these neurons are sampled randomly.

These empirically-grounded simulations demonstrate that modeling single-cell recordings and fMRI measurements as projections of the neural response space can reveal the dissimilarities between real neural patterns when the correlation distance is used as the RDM estimator. Moreover, they highlight the necessity and sufficience of removing the mean from the measured patterns —- an operation performed implicitly by the correlation distance—to uncover the close correspondence between these two measurement modalities. In contrast, estimating RDMs directly from raw patterns fails to achieve this close match.

## 3 Discussion

Our theoretical and simulation results promise that electrophysiological recordings and fMRI can give us good estimates of the neural-population representational geometry in certain scenarios. However, it is important to understand the assumptions these results depend on. Our encouraging results hold when measurement channels sample neurons linearly with weights drawn independently from some univariate distribution. This assumption is not in general realistic for either neural recordings or fMRI voxels. However, it provides an important idealized reference.

For fMRI, the non-negative independent-weight-linear-combination-sampling (IWLCS) model captures a best-case scenario in one sense and a worst-case scenario in another. It is a best-case scenario in the sense that all directions orthogonal to the all-1 direction of the neural response space are sampled equally and in that mean-removal enables the recovery of the original distances in expectation. It is a worst-case scenario in that each voxel samples a large number of independent dimensions of the neural response space, leading to extreme cancellation of tuning curves, which entails shrinkage of the geometry by a large factor in all dimensions orthogonal to the all-1 dimension. The non-negative IWLCS model is probably closer to reality for fMRI voxels if we assume that the sampled units are not single neurons but cortical columns of similarly tuned neurons. This reduces the number *N* of underlying representational dimensions and the number *k* of dimensions linearly combined by each voxel. The non-negative IWLCS model then correctly reflects that each voxel tends to average a less diverse set of tuning curves with subsets of the neurons that are correlated in their tuning, which entails less severe cancellation of tuning curves.

We now summarize the key insights of this study. Fig. 7 gives an overview. The measurement models we consider can be categorized into three main types:

a. **Random projections (Fig. 7a)**: We consider the theoretical scenario of sampling the neural activity as linear combinations with the weights drawn from a zero-mean Gaussian distribution. Under these conditions, the pairwise distances of measured activity patterns remain undistorted in expectation. Moreover a relatively small number of channels (e.g. 100) is sufficient for good estimates of the representational geometry. Unfortunately, this scenario does not hold for any real measurement modality.
b. **Electrophysiological recordings (Fig. 7b)**: We consider measurements where the individual activity of neurons is sampled invasively. Two potential scenarios arise:

- **Random sampling**: When recordings effectively represent a random sample of all neurons in the area, the measured representational geometry remains undistorted. Again, a small number of channels (e.g. 100) is sufficient for good estimates of the representational geometry. Pattern mean removal is not required or desirable in this scenario; it slightly reduces the accuracy of the geometry estimate as shown in Fig. 4.
- **Biased sampling**: Situations in which the neural selectivities are organized in a spatially structured manner, and the array only samples a fraction of this structure, present challenges. For instance, if an array is placed over a region that shows a particular selectivity, the measured representational geometry might fail to capture stimulus distinctions encoded in a different subset of the neural population of interest. As a result, the apparent geometry can be distorted in complex ways. Taking such distortions into account in inference on brain-computational models would require modeling the measurement-related biases and our uncertainty about them.
c. **fMRI measurements (Fig. 7c)**: We consider voxels that sample local averages of neural activity with non-negative weights. Depending on the spatial organization of the encoded information, outcomes can vary:

- **Code is spatially unstructured or structured at a single scale**: A code that has no spatial organization justifies the assumption that the channels sample with non-negative weights drawn independently from identical distributions. Structure at a single scale would hold if there were clusters of similarly tuned neurons (e.g. cortical columns), but these clusters were randomly arranged in the region. The latter situation is a sweet spot for fMRI: the clustering reduces signal cancellation due to averaging in voxels, yet the random arrangement of the clusters prevents complex distortions of the representational geometry (beyond overemphasis on the neural population mean). Whether the code is unstructured or structured at a single scale, two scenarios must be considered:

i. If the neural population mean is constant across conditions, then the measured representational geometry remains undistorted in expectation. Removing the mean from each the measured response pattern is not required (though it also will not hurt) for accurate reflection of the neural population representational geometry in the measured squared Euclidean distances.
ii. If the neural population mean varies across conditions, the measured squared Euclidean RDM is a linear combination of the original neuronal pattern RDM and the population-mean activation RDM. In this case, removing the mean from the measured patterns provides an undistorted estimate of the underlying representational geometry.
- **Code is spatially structured at multiple scales**: Codes that are structured at multiple scales will yield RDMs distorted in complex ways, reflecting the degrees to which different dimensions of the neural population code are cancelled by averaging inside voxels. For example, V1 is structured locally into orientation columns and globally into a retinotopic map. Orientation signals will be reduced substantially by averaging in voxels, whereas location signals will cancel much less because they reside at a coarser scale and are therefore more strongly reflected in fMRI patterns. Accounting for such complex distortions in evaluating brain-computational models requires models that predict the multi-scale spatial structure of the code, such that the distortions arising from locally-averaging voxels can be accounted for

**Figure 7:**
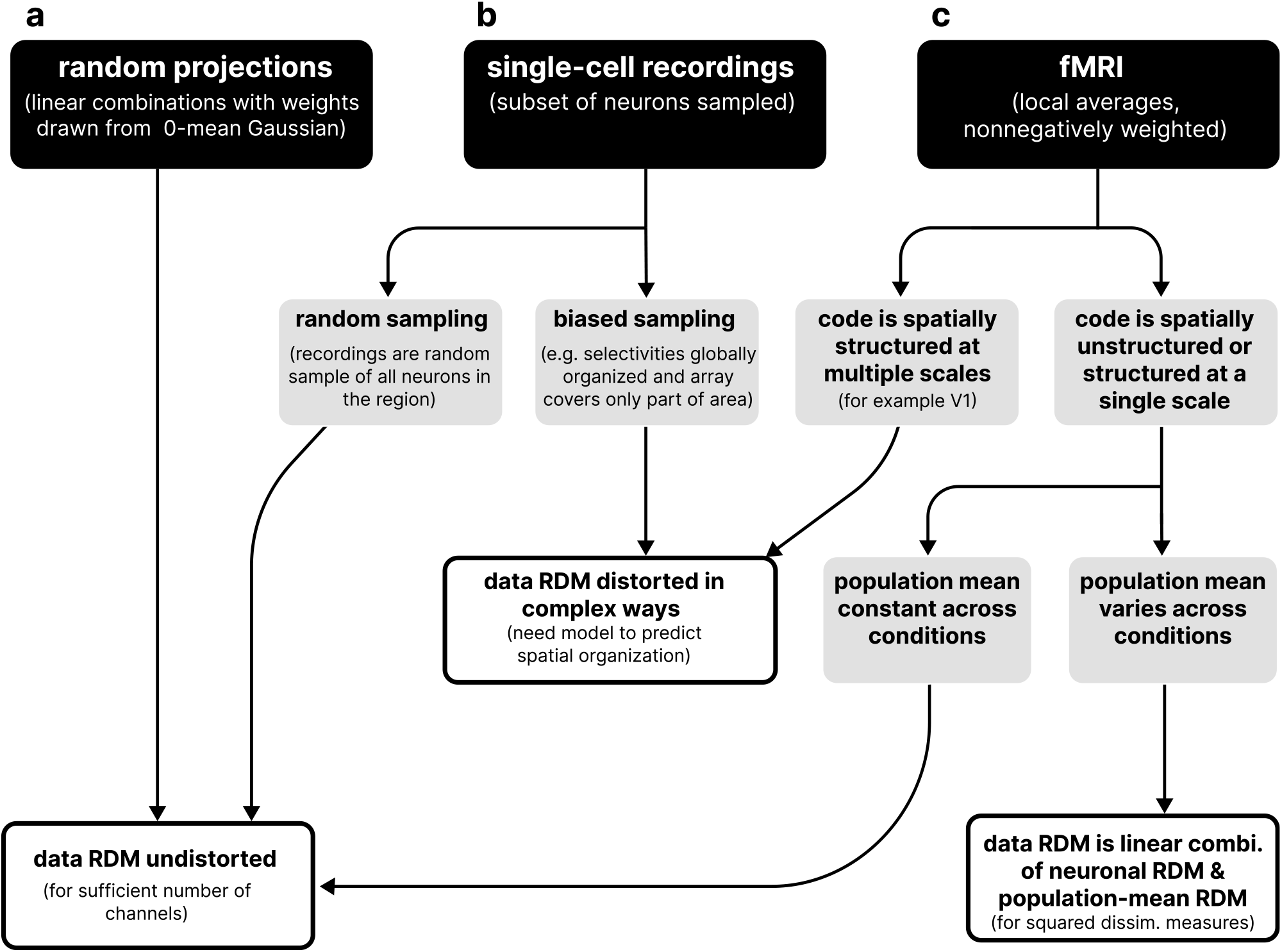
When do measured representational distances reflect the neural representational geometry? Three scenarios of brain-activity measurement conditions and their consequences for the resulting representational geometry. **a.** In the theoretical case of random projection sampling, the measured representational dissimilarities are an unbiased estimate of the neural pattern representational dissimilarities. **b.** For electrophysiological recordings, we consider two scenarios. In the first, the sampled cells are randomly selected from the underlying population, in which case the theoretical results and simulations in this study indicate that the apparent geometry remains undistorted. In the second scenario, where cells are spatially organized and measurements sample only a region of the population, the apparent representational geometry is expected to be distorted in complex ways, requiring separate modeling.. **c.** When sampling with fMRI voxels, if the underlying neural code is spatially structured at multiple scales, the geometry of the measured patterns will be distorted in complex ways that require modeling. In contrast, if the neural code is unstructured, two cases arise. First, if the mean activation across neurons is the same across all experimental conditions, the RDM derived from the measurements will provide an undistorted estimate of the neuronal RDM. Second, if the population mean varies across conditions, the apparent geometry will be linearly distorted along the population mean dimension. In this case, removing the mean activation from each measured pattern restores the underlying neural RDM in expectation.

### Multiple distortions affecting estimates of representational geometry

In this paper, we analyzed a particular source of distortion of estimates of the representational geometry that arises from the way measurement channels sample neural activity. It is important to note that the extent to which the measured geometry will be able to capture the true geometry is also affected by the fact that the measurements provide noisy estimates of the true activity patterns. A well-understood additional source of distortion that occurs when measuring representational geometries from brain activity data is the positive bias in the estimated distances between measured patterns. Consider the case where the neural patterns are identical (i.e. their true distance is zero). The measurement noise will cause errors in the pattern estimates, which will yield distances larger than 0. In general, high-dimensional measured patterns will tend to be further apart than the underlying true (i.e. noise free) patterns. This positive bias increases with the level of noise. This challenge has been investigated previously [45], and cross-validated distance estimators [28, 37, 3, 11, 9] have been introduced to remove it.

Our findings here demonstrate the overrepresentation of the population-mean dimension in measurements that average neural responses—a separate phenomenon independent of the noise. While mean removal can mitigate this, it will not mitigate the distortion effects induced by noise. In order to account for both of these sources of distortion, cross-validated distance estimators can be applied to pattern estimates after mean removal.

### Removing the mean from voxel patterns in fMRI measurements offers the benefits of the correlation distance while avoiding its drawbacks

The correlation distance between two patterns (1 *— r*, where *r* is the Pearson correlation coefficient) is proportional to the squared Euclidean distance computed after normalizing each pattern by subtracting its mean and dividing by its standard deviation (thus normalizing the mean to 0 and the variance to 1). The divisive pattern normalization introduces a notable pitfall: the resulting correlation distance does not reliably reflect linear discriminability. Consider two patterns corresponding to different conditions neither of which drives a significant response in our region of interest. The pattern estimates are dominated by noise and will be essentially uncorrelated (*r ≈* 0). Consequently, the correlation distance will be close to 1, suggesting that the patterns are highly dissimilar. However, a linear decoder would find that two conditions that drive no distinct responses are not decodable. If we remove the mean from fMRI pattern measurements, but do not perform variance normalization, we get the advantage of the correlation distance (correcting for the overrepresentation of the neural population mean dimension) while avoiding its pitfall (weak relationship to decodability).

## 4 Methods

### 4.1 Random geometry simulations

#### 4.1.1 Simulated activity patterns

In each simulation, we generated a random “ground truth” geometry of neural activity response patterns. We generated 100 data points derived from a Gaussian mixture in a five-dimensional space. These points were then embedded into a 1024-dimensional space via a random rotation. This resulted in the simulated activity of a population of 1024 neurons to 100 different experimental conditions. The proportion of the total response pattern variance attributed to the mean dimension averaged 25% across all simulations. We created a new random geometry for each new simulation.

#### 4.1.2 Measurement models

We sampled the simulated neural activity patterns using models that mimic certain properties of different brain-activity measurement modalities.

- **Zero-mean independent-weight-linear-combination-sampling (IWLCS) model.** This model samples a linear combination of the activity across the entire neuron population, with weights independently drawn from a standard Gaussian distribution. The resulting simulation approximates a Gaussian random projection, providing expected undistorted measurements. This model represents ideal sampling with random projections. It serves as a reference, rather than as a model of any actual measurement modality.
- **Non-negative IWLCS.** In this model, each channel samples a linear combination of the neural activity across the entire simulated neuron population. Weights are independently drawn from a truncated standard normal distribution (negative half set to zero), ensuring all weights are non-negative.
- **Sparse non-negative IWLCS model.** As in the previous model, each channel samples a linear combination from the neuron population. However, the weights are zero for a random 90% of the neurons, and drawn from a half-normal distribution for the remaining 10%.
- **Sparse zero-mean IWLCS model.** A sparse variation of the Gaussian model is also included, where 90% of the weights are zero.
- **Random subpopulation model.** This model emulates single-cell recording of brain activity by sampling the activity of a randomly selected subset of neurons in the population, with one simulated neuron represented per channel. Note that this model is not an IWLCS model, since the sampling axes are one-hot vectors, with exactly one neuronal weight equal to 1 and all other weights equal to 0. As the weights are not sampled identically and independently, this model does not satisfy the IWLCS assumptions.

#### 4.1.3 Representational geometry comparisons

We quantified the extent of similarity between the relationships among the neuronal patterns and those among the measured patterns by comparing their representational dissimilarity matrices (RDMs).

We computed the average Pearson correlation between pairs of RDMs across simulations. Six comparisons were performed in total. For each measurement model, we calculated the correlation between: a.) The RDMs of the squared Euclidean distances derived from simulated neuronal patterns (neuron RDM) and those derived from the measured patterns (measured RDM). b.) The one-dimensional RDM of absolute differences between the mean activations in the simulated neural population (neuron mean RDM) and the one-dimensional RDM of absolute differences between the means across the measurement channels (measured mean RDM). c.) The neuron RDM and the RDM of squared Euclidean distances from the simulated measured patterns, after removing the mean measurement from each pattern (mean-removed measured RDM). d.) The RDM of squared Euclidean distances from the simulated neural response patterns, after removing the population mean activation from each pattern, and the mean-removed measured RDM. e.) The neuron RDM and the RDM consisting of correlation distances between the simulated measured patterns (measured correlation RDM). f.) The RDM of correlation distances from the simulated neuronal patterns (neuron correlation RDM) and the measured correlation RDM.

### 4.2 Simulations based on neural data

#### Empirical data

The neural recording dataset underpinning our simulations is from the study [23] and has also been used in [31]. The latter study compared the representational geometry measured with fMRI in the human ventral temporal cortex to the representational geometry estimated from neural recordings in macaque inferior temporal cortex (IT) in response to images of various object categories. We used the 92 *×* 92 representational dissimilarity matrix for macaque IT used in [31] as a basis for our simulations.

#### Ground-truth neural pattern generation

Using the observed representational dissimilarities from monkey IT, we generated embeddings of neural activation patterns in a 1000-dimensional simulated neural response space. To do so, we converted the monkey IT RDM (correlation distances between patterns) into a positive definite linear kernel (similarity) matrix *K* by double centering and multiplying by -0.5 [10]. We then used the Cholesky decomposition of the kernel matrix (*K* = *LL^>^*) to sample a random stimulus-response matrix conforming exactly to the kernel matrix. To do this, we first sampled a random matrix *U* from a standard normal distribution and normalized it such that its rows had zero mean and unit variance, ensuring no prior structure was imposed. We then multiplied *U* by the Cholesky factor *L*, effectively mapping the random matrix into the space defined by the kernel *K*. Because of the equivalence between linear kernel matrices and squared Euclidean distance matrices, this procedure ensures that the squared Euclidean distances between the patterns in the stimulus-response matrix match the ground-truth RDM. This method for simulating neural response patterns is implemented in the Python RSA toolbox (<u>github.com/rsagroup/rsatoolbox</u>, [43]).

The resulting patterns all have the same mean activation. In order to illustrate the distortion that sampling with simulated fMRI voxels creates in the apparent geometry, it is necessary that there be some variation among these means (otherwise, the term in Eq. 1 would be zero for every pair, and the distance estimates would be undistorted in expectation). To introduce variation along the mean, we added a random mean to each of the 92 patterns. The squared Euclidean distances between these patterns are consistent with the Kriegeskorte et al. study (after mean-removal) and served as our simulated ground-truth neural population for subsequent analysis.

## 5 Author contributions

N.K. conceived the study. V.B.B. ran simulations and analyzed data. V.B.B. and N.K. interpreted the theoretical and simulation results. V.B.B. and N.K. wrote and edited the manuscript.

