## Supplemental material for "When do measured representational distances reflect the neural representational geometry?"

### 7 Supplemental material

#### 7.1 Measured dissimilarities are unbiased for zero-mean neural sampling weights or equal-mean neural patterns: Proof of Equation 1

Let $\mathbf{x}, \mathbf{y} \in \mathbb{R}^N$ be response patterns in neuronal space. Let $\{\mathbf{w}_1, ..., \mathbf{w}_M\}$ be random vectors whose entries $w_{ij}$ are drawn i.i.d. from a given univariate probability distribution $p(w)$.

Let $\mathbf{W}$ be the $M \times N$ matrix whose rows are $\{\mathbf{w}_1, ..., \mathbf{w}_M\}$. Consider the measurement transformation $\pi : \mathbb{R}^N \to \mathbb{R}^M$ where $\pi(\mathbf{x}) = \mathbf{W} \cdot \mathbf{x}$ is the measured response pattern (e.g. voxel pattern) whose entries are linear combinations of the neural responses. Then for any pair of neural response patterns $\mathbf{x}, \mathbf{y}$, the expected value of the squared Euclidean distance of their measured patterns $\pi(\mathbf{x}), \pi(\mathbf{y})$ (scaled by a factor of $\frac{1}{M}$) will be:

$$
\begin{aligned}
\mathbb{E}\left[\frac{1}{M}\|\pi(\mathbf{x}) - \pi(\mathbf{y})\|^2\right] &= \mathop{\mathbb{E}}_{w_{ij}\sim p(w)}\left[\frac{1}{M}\sum_{i=1}^{M}\Big(\sum_{j=1}^{N} w_{ij}x_j - \sum_{i=1}^{N} w_{ij}y_j\Big)^2\right] && (6)\\
&= \mathop{\mathbb{E}}_{w_{ij}\sim p(w)}\left[\frac{1}{M}\sum_{i=1}^{M}\Big(\sum_{j=1}^{N} w_{ij}(x_j - y_j)\Big)^2\right] && (7)\\
&= \mathop{\mathbb{E}}_{w_{j}\sim p(w)}\left[\Big(\sum_{j=1}^{N} w_{j}(x_j - y_j)\Big)^2\right] && (8)\\
&= \mathop{\mathbb{E}}_{w_{j}\sim p(w)}\left[\sum_{j=1}^{N}\sum_{k=1}^{N} w_j w_k (x_j - y_j)(x_k - y_k)\right] && (9)\\
&= \mathbb{E}\left[\sum_j w_j^2 (x_j - y_j)^2\right] + \mathbb{E}\left[\sum_j \sum_{k\neq j} w_j w_k (x_j - y_j)(x_k - y_k)\right] && (10)\\
&= \Big(\sum_j \mathbb{E}[w_j^2]\cdot(x_j - y_j)^2\Big) + \Big(\sum_j\sum_{k\neq j}\mathbb{E}[w_j w_k]\cdot(x_j - y_j)(x_k - y_k)\Big) && (11)\\
&= \mathbb{E}\left[w^2\right]\cdot\sum_j (x_j - y_j)^2 + \mathbb{E}\left[w\right]^2\cdot\sum_j\sum_{k\neq j}(x_j - y_j)(x_k - y_k) && (12)\\
&= \mathbb{E}\left[w^2\right]\cdot\|\mathbf{x} - \mathbf{y}\|^2 + \mathbb{E}\left[w\right]^2\cdot\left[\sum_{j,k}(x_j - y_j)(x_k - y_k) - \sum_j (x_j - y_j)^2\right] && (13)\\
&= \Big(\mathbb{E}\left[w^2\right] - \mathbb{E}\left[w\right]^2\Big)\cdot\|\mathbf{x} - \mathbf{y}\|^2 + \mathbb{E}\left[w\right]^2\cdot\sum_{j,k}(x_j - y_j)(x_k - y_k) && (14)
\end{aligned}
$$

$$= \left(\mathbb{E}\left[w^2\right] - \mathbb{E}\left[w\right]^2\right) \cdot \|\mathbf{x}-\mathbf{y}\|^2 + \mathbb{E}\left[w\right]^2 \cdot \sum_j \left[(x_j - y_j) \cdot \textstyle\sum_k (x_k - y_k)\right] \tag{15}$$

$$= \left(\mathbb{E}\left[w^2\right] - \mathbb{E}\left[w\right]^2\right) \cdot \|\mathbf{x}-\mathbf{y}\|^2 + \mathbb{E}\left[w\right]^2 \cdot \sum_k (x_k - y_k) \cdot \sum_j (x_j - y_j) \tag{16}$$

$$= \left(\mathbb{E}\left[w^2\right] - \mathbb{E}\left[w\right]^2\right) \cdot \|\mathbf{x}-\mathbf{y}\|^2 + \mathbb{E}\left[w\right]^2 \cdot \left(\sum_j (x_j - y_j)\right)^2 \tag{17}$$

$$= \left(\mathbb{E}\left[w^2\right] - \mathbb{E}\left[w\right]^2\right) \cdot \|\mathbf{x}-\mathbf{y}\|^2 + \mathbb{E}\left[w\right]^2 \cdot \left(\sum_j x_j - \sum_j y_j\right)^2 \tag{18}$$

$$= \mathrm{Var}[w] \cdot \|\mathbf{x}-\mathbf{y}\|^2 + \mathbb{E}\left[w\right]^2 \cdot (\mathbb{1}^\top \mathbf{x} - \mathbb{1}^\top \mathbf{y})^2 \tag{19}$$

Note that if the distribution of $w$ has mean 0 or patterns $\mathbf{x}$ and $\mathbf{y}$ have the same mean activity, the final expression reduces to:

$$\mathrm{Var}[w] \cdot \|\mathbf{x}-\mathbf{y}\|^2$$

If, additionally, the variance is 1, we get the squared Euclidean distance of $\mathbf{x}, \mathbf{y}$ in neuron space. That is, if $w_{ij}$ are drawn i.i.d. from a distribution with $\mathbb{E}(w_{ij}) = 0$ and $\mathrm{Var}(w_{ij}) = 1$, then the squared Euclidean distances computed from the measurement channels provide an unbiased estimate of the squared Euclidean distances of the neural population.

#### 7.2 The apparent representational geometry is linearly stretched along the all-1 direction of the neural response space

##### Linear stretch factor

Consider an arbitrary pair of neural response patterns $\mathbf{x}$ and $\mathbf{y}$ in $\mathbb{R}^N$. The squared distance between these vectors can be decomposed as follows:

$$\|\mathbf{x}-\mathbf{y}\|^2 = \|\mathbf{x}-\mathbf{y}\|^2_{\text{along}} + \|\mathbf{x}-\mathbf{y}\|^2_{\text{ortho}} \tag{20}$$

where $\|\mathbf{x}-\mathbf{y}\|^2_{\text{along}}$ represents the component of the distance that lies along the all-one vector ($\mathbb{1}$), and $\|\mathbf{x}-\mathbf{y}\|^2_{\text{ortho}}$ represents the component of the distance orthogonal to the all-one vector:

$$||\mathbf{x}-\mathbf{y}||^2_{\text{along}} = \|\mathrm{proj}_{\mathbb{1}}(\mathbf{x}-\mathbf{y})\|^2 = \frac{1}{N}(\mathbb{1}^\top \mathbf{x} - \mathbb{1}^\top \mathbf{y})^2 \tag{21}$$

$$||\mathbf{x}-\mathbf{y}||^2_{\text{ortho}} = \sum (x_i - y_i)^2 - \frac{1}{N}(\mathbb{1}^\top \mathbf{x} - \mathbb{1}^\top \mathbf{y})^2 \tag{22}$$

Thus, by replacing this sum into Eq. 1, we can observe how these two components are represented in the measurements under the i.i.d sampling model defined in section 2.1.1.

Note that $(\mathbb{1}^\top \mathbf{x} - \mathbb{1}^\top \mathbf{y})^2 = N \cdot ||\mathbf{x} - \mathbf{y}||^2_{\text{along}}$. So,

$$
\begin{aligned}
\mathbb{E}\left[\frac{1}{M}\|\pi(\mathbf{x}) - \pi(\mathbf{y})\|^2\right] &= \text{Var}[w] \cdot \|\mathbf{x} - \mathbf{y}\|^2 + \mathbb{E}[w]^2 \cdot (\mathbb{1}^\top \mathbf{x} - \mathbb{1}^\top \mathbf{y})^2 \\
&= \text{Var}[w]\left[\|\mathbf{x} - \mathbf{y}\|^2_{\text{along}} + \|\mathbf{x} - \mathbf{y}\|^2_{\text{ortho}}\right] + \mathbb{E}[w]^2 \cdot N \cdot \|\mathbf{x} - \mathbf{y}\|^2_{\text{along}} \\
&= (\text{Var}[w]) \cdot \|\mathbf{x} - \mathbf{y}\|^2_{\text{ortho}} + (\text{Var}[w] + \mathbb{E}[w]^2 \cdot N) \cdot \|\mathbf{x} - \mathbf{y}\|^2_{\text{along}}
\end{aligned}
\tag{23}
$$

The ratio of the factors that accompany each of the terms in the decomposition, namely, $\frac{\text{Var}[w]}{\text{Var}[w] + \mathbb{E}[w]^2 \cdot N}$ provide a measure of the factor by which the representational geometry is shrunk along the orthogonal dimensions (or stretched along the all-1 dimension) by the measurement.

#### 7.3 Dissimilarities among the mean-removed measured patterns provide unbiased estimates of neural dissimilarities.

##### 7.3.1 Proof of Equation 4

Recall Eq. 4:

$$\bar{\mathbf{v}} = \mathbf{C} \cdot \mathbf{v} = \mathbf{C} \cdot \mathbf{W}\mathbf{x} = \bar{\mathbf{W}}\mathbf{x}$$

For each measured pattern $\mathbf{v} = \pi(\mathbf{x})$, we compute the mean-removed measured pattern $\bar{\pi}(\mathbf{x})$ as follows:

$$\bar{\pi}(\mathbf{x}) = \pi(\mathbf{x}) - \mathbb{1}\frac{1}{M}\mathbb{1}^\top \pi(\mathbf{x}) = \pi(\mathbf{x}) - \frac{\mathbb{1}\mathbb{1}^\top}{M}\pi(\mathbf{x}) \tag{24}$$

Substituting the definition of $\pi$ (Eq. 2), we have:

$$
\begin{aligned}
\bar{\pi}(\mathbf{x}) &= \mathbf{W}\mathbf{x} - \frac{\mathbb{1}\mathbb{1}^\top}{M}\mathbf{W}\mathbf{x} \\
&= \left(\mathbf{I}_M - \frac{\mathbb{1}\mathbb{1}^\top}{M}\right) \cdot \mathbf{W}\mathbf{x} \\
&= \bar{\mathbf{W}}\mathbf{x},
\end{aligned}
\tag{25}
$$

where

$$\bar{\mathbf{W}} = \left(\mathbf{I}_M - \frac{\mathbb{1}\mathbb{1}^\top}{M}\right) \cdot \mathbf{W}$$

$\bar{\mathbf{W}}$ is the mean-removed weights matrix, where each column of $\mathbf{W}$ has been separately centered. So we see that centering each measured pattern is equivalent to centering each column of the weights matrix.

For intuition, the key thing to appreciate is that removing the mean separately from each measured pattern is equivalent to centering each *column* of $\mathbf{W}$. What does this do to the ensemble of weight vectors $\mathbf{w}_i = [w_{i1}, ..w_{iN}]$ (the *rows* of $\mathbf{W}$) in the neural population response space? It rigidly centers these vectors on the origin of neural population response space. This intuition makes it clear, that the neural-population-mean dimension is now sampled just like all other neural-response-space dimensions.

###### 7.3.2 Proof of equation 5

Eq. 5 states that subtracting the mean from each of the measured response patterns $\pi(\mathbf{x})$ and $\pi(\mathbf{y})$ to obtain the centered measured response patterns $\bar{\pi}(\mathbf{x})$ and $\bar{\pi}(\mathbf{x})$ yields an unbiased estimate of the representational distance between the underlying neural patterns $\mathbf{x}$ and $\mathbf{y}$, up to a scaling factor:

$$E\left[\frac{1}{M}\left\|\bar{\pi}(\mathbf{y})-\bar{\pi}(\mathbf{x})\right\|^2\right]=\frac{M-1}{M}\operatorname{Var}[w]\left\|\mathbf{y}-\mathbf{x}\right\|^2$$

Thus, the expected value of the squared Euclidean distance between the centered measured response patterns is $(M-1)\cdot\operatorname{Var}[w]$ times the squared Euclidean distance $\|\mathbf{y}-\mathbf{x}\|^2$ between the neural response patterns.

To see why this is so, first note that for each new weight $\bar{w}_{ij}=w_{ij}-\frac{1}{M}\sum_{k=1}^{M}w_{kj}$.

$$\begin{aligned}
\mathbb{E}[\bar{w}_{ij}\cdot\bar{w}_{ij}] &= \mathbb{E}\left[(w_{ij}-\frac{1}{M}\sum_{p=1}^{M}w_{pj})(w_{ij}-\frac{1}{M}\sum_{p=1}^{M}w_{pj})\right]\\
&= \mathbb{E}\left[w_{ij}w_{ij}-w_{ij}\frac{1}{M}\sum_{p}w_{pj}-w_{ij}\frac{1}{M}\sum_{p}w_{pj}+\frac{1}{M^2}\left(\sum_{p}w_{pj}\right)^2\right]\\
&= \mathbb{E}[w_{ij}^2]-\frac{1}{M}\sum_{p=1}^{M}\mathbb{E}[w_{ij}w_{pj}]-\frac{1}{M}\sum_{p=1}^{M}\mathbb{E}[w_{ij}w_{pj}]+\frac{1}{M^2}\left(\sum_{p=1}^{M}\mathbb{E}[w^2]+\sum_{p=1}^{M}\sum_{l\neq p}E[w_{pj}w_{lj}]\right)\\
&= \mathbb{E}[w^2]-\frac{2}{M}\sum_{p=1}^{M}\mathbb{E}[w_{ij}w_{pj}]+\frac{1}{M^2}\left(\sum_{p=1}^{M}\mathbb{E}[w^2]+\sum_{p=1}^{M}\sum_{l\neq p}E[w_{pj}w_{lj}]\right)\\
&= \mathbb{E}[w^2]-\frac{2}{M}\left(\mathbb{E}[w^2]+M-1\,\mathbb{E}[w]^2\right)+\frac{1}{M^2}\left(M\,\mathbb{E}[w^2]+M(M-1)E[w]^2\right)\\
&= \mathbb{E}[w^2]\cdot\left(1-\frac{1}{M}\right)-\frac{2}{M}\left(M-1\,\mathbb{E}[w]^2\right)+\frac{1}{M^2}\left(M(M-1)E[w]^2\right)\\
&= \mathbb{E}[w^2]\cdot\left(1-\frac{1}{M}\right)-2\cdot\left(\frac{M-1}{M}\mathbb{E}[w]^2\right)+\frac{1}{M^2}\left(M(M-1)E[w]^2\right)\\
&= \mathbb{E}[w^2]\cdot\left(1-\frac{1}{M}\right)-2\cdot\left(\frac{M-1}{M}\mathbb{E}[w]^2\right)+\frac{M-1}{M}\left(E[w]^2\right)\\
&= \mathbb{E}[w^2]\cdot\frac{M-1}{M}-\mathbb{E}[w]^2\cdot\frac{M-1}{M}\\
&= \frac{M-1}{M}\operatorname{Var}[w]
\end{aligned} \tag{26}$$

Now,

$$
\begin{aligned}
\mathbb{E}\left[\frac{1}{M}\left\|\bar{\pi}(\mathbf{y})-\bar{\pi}(\mathbf{x})\right\|^2\right] &= \mathbb{E}\left[\frac{1}{M}\left\|\bar{\mathbf{W}}\mathbf{y}-\bar{\mathbf{W}}\mathbf{x}\right\|^2\right] \\
&= \mathbb{E}\left[\frac{1}{M}\sum_{i=1}^{M}\left(\sum_{j=1}^{N}\bar{w}_{ij}(y_j-x_j)\right)^2\right] \\
&= \frac{1}{M}\sum_{i=1}^{M}\mathbb{E}\left[\sum_{j}\sum_{k\neq j}\bar{w}_{ij}\bar{w}_{ik}(y_j-x_j)(y_k-x_k)+\sum_{j=1}^{N}\bar{w}_{ij}^2(y_j-x_j)^2\right] \\
&= \frac{1}{M}\sum_{i=1}^{M}\left(\mathbb{E}\left[\sum_{j}\sum_{k\neq j}\bar{w}_{ij}\bar{w}_{ik}(y_j-x_j)(y_k-x_k)\right]+\mathbb{E}[\bar{w}^2]\cdot\|\mathbf{y}-\mathbf{x}\|^2\right) \\
&= \frac{1}{M}\sum_{i=1}^{M}\left(\sum_{j}\sum_{k\neq j}\mathbb{E}[\bar{w}_{ij}\bar{w}_{ik}](y_j-x_j)(y_k-x_k)+\mathbb{E}[\bar{w}^2]\cdot\|\mathbf{y}-\mathbf{x}\|^2\right) \\
&= \frac{1}{M}\sum_{i=1}^{M}\left(\sum_{j}\sum_{k\neq j}\mathbb{E}[\bar{w}_{ij}]\,\mathbb{E}[\bar{w}_{ik}](y_j-x_j)(y_k-x_k)+\mathbb{E}[\bar{w}^2]\cdot\|\mathbf{y}-\mathbf{x}\|^2\right) \\
&= \frac{1}{M}\sum_{i=1}^{M}\left(\sum_{j}\sum_{k\neq j}\mathbb{E}[\bar{w}]^2(y_j-x_j)(y_k-x_k)+\mathbb{E}[\bar{w}^2]\cdot\|\mathbf{y}-\mathbf{x}\|^2\right) \\
&= \frac{1}{M}\sum_{i=1}^{M}\left(\sum_{j}\mathbb{E}[\bar{w}]^2(y_j-x_j)^2+\mathbb{E}[\bar{w}^2]\cdot\|\mathbf{y}-\mathbf{x}\|^2\right) \\
&= \frac{1}{M}\cdot M\cdot\mathbb{E}[\bar{w}^2]\cdot\|\mathbf{y}-\mathbf{x}\|^2 \\
&= \frac{M-1}{M}\operatorname{Var}[w]\,\|\mathbf{y}-\mathbf{x}\|^2
\end{aligned} \tag{27}
$$

##### 7.4 The squared Euclidean distance computed after mean removal and variance normalization equals twice the Pearson correlation distance

Let $\mathbf{x}$ and $\mathbf{y}$ be two random variables in $\mathbb{R}^N$. Let $\bar{x} = \frac{1}{N}\sum_{i=1}^{N} x_i$ and $s_x = \sqrt{\frac{1}{N}\sum(x_i-\bar{x})^2}$, and analogously for $\bar{y}$ and $s_y$. The Pearson correlation coefficient between $x$ and $y$ is defined by:

$$
r_{xy} = \frac{1}{N}\sum_{i=1}^{N}\left(\frac{x_i-\bar{x}}{s_x}\right)\left(\frac{y_i-\bar{y}}{s_y}\right)
$$

Now let's consider the squared Euclidean distance between $\mathbf{x}$ and $\mathbf{y}$ after removing their mean and normalizing to unit variance:

$$\begin{aligned}
\frac{1}{N}d^2\left(\frac{\mathbf{x}-\bar{x}}{s_x}-\frac{\mathbf{y}-\bar{y}}{s_y}\right) &= \frac{1}{N}||\frac{\mathbf{x}-\bar{x}}{s_x}-\frac{\mathbf{y}-\bar{y}}{s_y}||^2 \\
&= \frac{1}{N}\left(\sum_{i=1}^{N}\left(\frac{x_i-\bar{x}}{s_x}-\frac{y_i-\bar{y}}{s_y}\right)^2\right) \\
&= \frac{1}{N}\left(\sum_{i=1}^{N}\left(\frac{x_i-\bar{x}}{s_x}\right)^2-2\sum_{i=1}^{N}\left(\frac{x_i-\bar{x}}{s_x}\right)\left(\frac{y_i-\bar{y}}{s_y}\right)+\sum_{i=1}^{N}\left(\frac{y_i-\bar{y}}{s_y}\right)^2\right) \\
&= \frac{1}{N}\left(\frac{1}{s_x^2}Ns_x^2-2\sum_{i=1}^{N}\left(\frac{x_i-\bar{x}}{s_x}\right)\left(\frac{y_i-\bar{y}}{s_y}\right)+\frac{1}{s_x^2}Ns_x^2\right) \\
&= \frac{1}{N}\left(2N-2\sum_{i=1}^{N}\left(\frac{x_i-\bar{x}}{s_x}\right)\left(\frac{y_i-\bar{y}}{s_y}\right)\right) \\
&= \frac{1}{N}\left(2N-2\cdot Nr_{xy}\right) \\
&= 2(1-r_{xy})
\end{aligned}$$

#### 7.5 Linear shrink factor of the apparent representational geometry due to voxel sampling (linear model)

**Distance distortion decomposed along and orthogonal to all-1 dimension of the neural response space**

Consider an arbitrary pair of neural respose patterns $\mathbf{x}$ and $\mathbf{y}$ in $\mathbb{R}^N$. The squared distance between these vectors can be decomposed as follows:

$$\|\mathbf{x}-\mathbf{y}\|^2 = \|\mathbf{x}-\mathbf{y}\|^2_{\text{along}} + \|\mathbf{x}-\mathbf{y}\|^2_{\text{ortho}} \tag{28}$$

where $\|\mathbf{x}-\mathbf{y}\|^2_{\text{along}}$ represents the component of the distance that lies along the all-one vector, and $\|\mathbf{x}-\mathbf{y}\|^2_{\text{ortho}}$ represents the component of the distance orthogonal to the all-one vector:

$$||\mathbf{x}-\mathbf{y}||^2_{\text{along}} = \|\text{proj}_{\mathbb{1}}(\mathbf{x}-\mathbf{y})\|^2 = \frac{1}{N}(\mathbb{1}^\top\mathbf{x}-\mathbb{1}^\top\mathbf{y})^2 \tag{29}$$

$$||\mathbf{x}-\mathbf{y}||^2_{\text{ortho}} = \sum(x_i-y_i)^2 - \frac{1}{N}(\mathbb{1}^\top\mathbf{x}-\mathbb{1}^\top\mathbf{y})^2 \tag{30}$$

Thus, by replacing this sum into equation 1, we can observe how these two components are represented in the measurements under the i.i.d sampling model defined in section 2.1.1:

(31)

Note that $(\mathbb{1}^\top \mathbf{x} - \mathbb{1}^\top \mathbf{y})^2 = N \cdot ||\mathbf{x} - \mathbf{y}||^2_{\text{along}}$

So that Eq. 7.5 becomes:

$$\begin{aligned}
\mathbb{E}\left[\frac{1}{M}\|\pi(\mathbf{x}) - \pi(\mathbf{y})\|^2\right] &= \text{Var}[w] \cdot \|\mathbf{x} - \mathbf{y}\|^2 + \mathbb{E}[w]^2 \cdot (\mathbb{1}^\top \mathbf{x} - \mathbb{1}^\top \mathbf{y})^2 \\
&= \text{Var}[w]\left[\|\mathbf{x} - \mathbf{y}\|^2_{\text{along}} + \|\mathbf{x} - \mathbf{y}\|^2_{\text{ortho}}\right] + \mathbb{E}[w]^2 \cdot N \cdot \|\mathbf{x} - \mathbf{y}\|^2_{\text{along}} \\
&= (\text{Var}[w]) \cdot \|\mathbf{x} - \mathbf{y}\|^2_{\text{ortho}} + (\text{Var}[w] + \mathbb{E}[w]^2 \cdot N) \cdot \|\mathbf{x} - \mathbf{y}\|^2_{\text{along}}
\end{aligned} \tag{32}$$

The ratio of the factors that accompany each of the terms in the decomposition, namely, $\frac{\text{Var}[w]}{\text{Var}[w] + \mathbb{E}[w]^2 \cdot N}$ will give us a measure of the factor by which the representational geometry is shrunk along the orthogonal dimensions (or stretched along the all-1 dimension) by the measurement.

#### 7.6 Geometry stretch factor for voxels that average $k$ neurons

We now consider how the representational geometry is distorted when each voxel samples a random subset of neurons. If each voxel sampled exactly $k$ neurons by averaging them, the sampling weights would not be independent and so Eq. 1 would not hold. However, it is unlikely that each voxel samples exactly $k$ neurons. We consider a sampling model, in which voxels sample $k$ neurons on average, each with an equal weight of $1/k$. The weights, thus, take values 0 and $1/k$ and are drawn independently.

Let the weights $w_{ij}$ be i.i.d. random variables such that:

$$w_{ij} = \begin{cases} \frac{1}{k} & \text{with probability } p = \frac{k}{N}, \\ 0 & \text{with probability } p = 1 - \frac{k}{N}. \end{cases}$$

The expected number of neurons sampled by a voxel then is $k/N \cdot N = k$. The expected value of the weights is:

$$\mathbb{E}[w_{ij}] = \frac{1}{k} \cdot \frac{k}{N} + 0 \cdot \left(1 - \frac{k}{N}\right) = \frac{1}{N}$$

The expected value of the squared weights is:

$$\mathbb{E}[w_{ij}^2] = \left(\frac{1}{k}\right)^2 \cdot \frac{k}{N} + 0^2 \cdot \left(1 - \frac{k}{N}\right) = \frac{1}{kN}$$

The variance of the weights is:

$$\text{Var}(w_{ij}) = \mathbb{E}[w_{ij}^2] - \mathbb{E}[w_{ij}]^2 = \frac{1}{kN} - \frac{1}{N^2} = \frac{N-k}{kN^2}$$

Entering the expected value of the weights $\mathbb{E}[w]$ and variance of the weights $\text{Var}[w]$ for this voxel-sampling model into Eq. 3, we get the shrink factor this model predicts:

$$\begin{aligned} f_{shrink} &= \left[\frac{\text{Var}[w]}{\text{Var}[w] + \mathbb{E}[w]^2 \cdot N}\right]^{\frac{1}{2}} = \left[\frac{\frac{N-k}{kN^2}}{\frac{N-k}{kN^2} + \frac{1}{N^2} \cdot N}\right]^{\frac{1}{2}} \\ &= \sqrt{\frac{N-k}{N-k+kN}} = \frac{1}{\sqrt{1 + \frac{kN}{N-k}}} \end{aligned} \tag{33}$$

In this case, it is more intuitive to consider the reciprocal of the shrink factor, which we call the stretch factor:

$$f_{stretch} = \sqrt{1 + \frac{kN}{N-k}} \tag{34}$$

Note that for $k = 1$, the stretch factor is already slightly bigger than $\sqrt{2}$ for any $N > 0$. This is a case where the i.i.d. sampling model we consider here deviates from the model where each voxel averages exactly $k$ neurons. The latter model, for $k = 1$, coincides with the subpopulation sampling model and has stretch factor $f_{stretch} = 1.0$. For substantially larger $k$, however, the i.i.d. model and the $k$-neuron-average model have similar stretch factors.

When sampling cortical areas with fMRI voxels, there are always millions of neurons in the population of interest and many thousands of neurons in each voxel. We may have hundreds of voxels sampling the cortical area. So realistically, $N$ will be huge and $k$ will be quite large but orders of magnitude smaller than $N$. In this $1 \ll k \ll N$ regime, the stretch factor for the Euclidean distances is close to $\sqrt{k}$: $f_{stretch} \approx \sqrt{k+1}$. (Since $N - k \approx N$, so $\frac{kN}{N-k}$ in eq 34 becomes very close to $k$).

This suggests that representational geometries estimated from 2-mm isotropic fMRI voxel patterns using the Euclidean distances without removing the mean will be distorted by huge factors. The number of neurons per voxel will be tens or hundreds of thousands. Assuming optimistically $k = 10,000$, the representational geometry would be stretched by factor $f_{stretch} = \sqrt{k} = 100$. Note, however, that the model assumes that each voxel samples a random subset of neurons. If we, perhaps more realistically, assume that each voxel randomly samples a smaller number of coarser-scale columns, results will differ. If each voxel sampled $k = 9$ columns, the representational geometry would be stretched by factor $f_{stretch} = \sqrt{k} = 3$.

The random sampling model may also be overly conservative in that real fMRI voxels are localized so as to sample disjoint spatial regions. Although a real fMRI voxel may sample somewhat overlapping sets of neurons due to mixing effects in the vasculature and at the image-reconstruction stage, the overlap may not be as severe as for independent sampling.
